# Effects of physical, chemical, and biological ageing on the mineralization of pine wood biochar by a *Streptomyces* isolate

**DOI:** 10.1101/2021.02.10.430652

**Authors:** Nayela Zeba, Timothy D. Berry, Thea L. Whitman

**Affiliations:** Department of Soil Science, University of Wisconsin-Madison, Madison, WI, 53703, USA

**Keywords:** Aging, Biochar C mineralization, FT-IR, Surface oxidation, O/C ratio, pyrogenic C, PyC, pyrogenic organic matter, PyOM

## Abstract

If biochar is to be used for carbon (C) management, we must understand how ageing affects biochar C mineralization. Here, we incubated aged and unaged eastern white pine wood biochar produced at 350 and 550 °C with a *Streptomyces* isolate, a putative biochar-decomposing microbe. Ageing was simulated via exposure to (a) alternating freeze-thaw and wet-dry cycles (physical ageing), (b) concentrated hydrogen peroxide (chemical ageing) and (c) nutrients and microorganisms (biological ageing). Elemental composition and surface chemistry (Fourier Transform Infrared spectroscopy) of biochar samples were compared before and after ageing. Ageing significantly increased biochar C mineralization in the case of physically aged 350 °C biochar (p < 0.001). Among 350 °C biochars, biochar C mineralization was positively correlated with an increase in O/C ratio (R^2^ = 0.78) and O-containing functional groups (R^2^ = 0.73) post-ageing, suggesting that surface oxidation during ageing enhanced biochar degradation by the isolate. However, in the case of 550 °C biochar, ageing did not result in a significant change in biochar C mineralization (p > 0.05), likely due to lower surface oxidation and high condensed aromatic C content. These results have implications for the use of biochar for long term C storage in soils.

**Synopsis:** This study highlights the impact of ageing on the microbial mineralization of biochar, which can affect its long-term C storage capacity.

**TOC Graphic:** 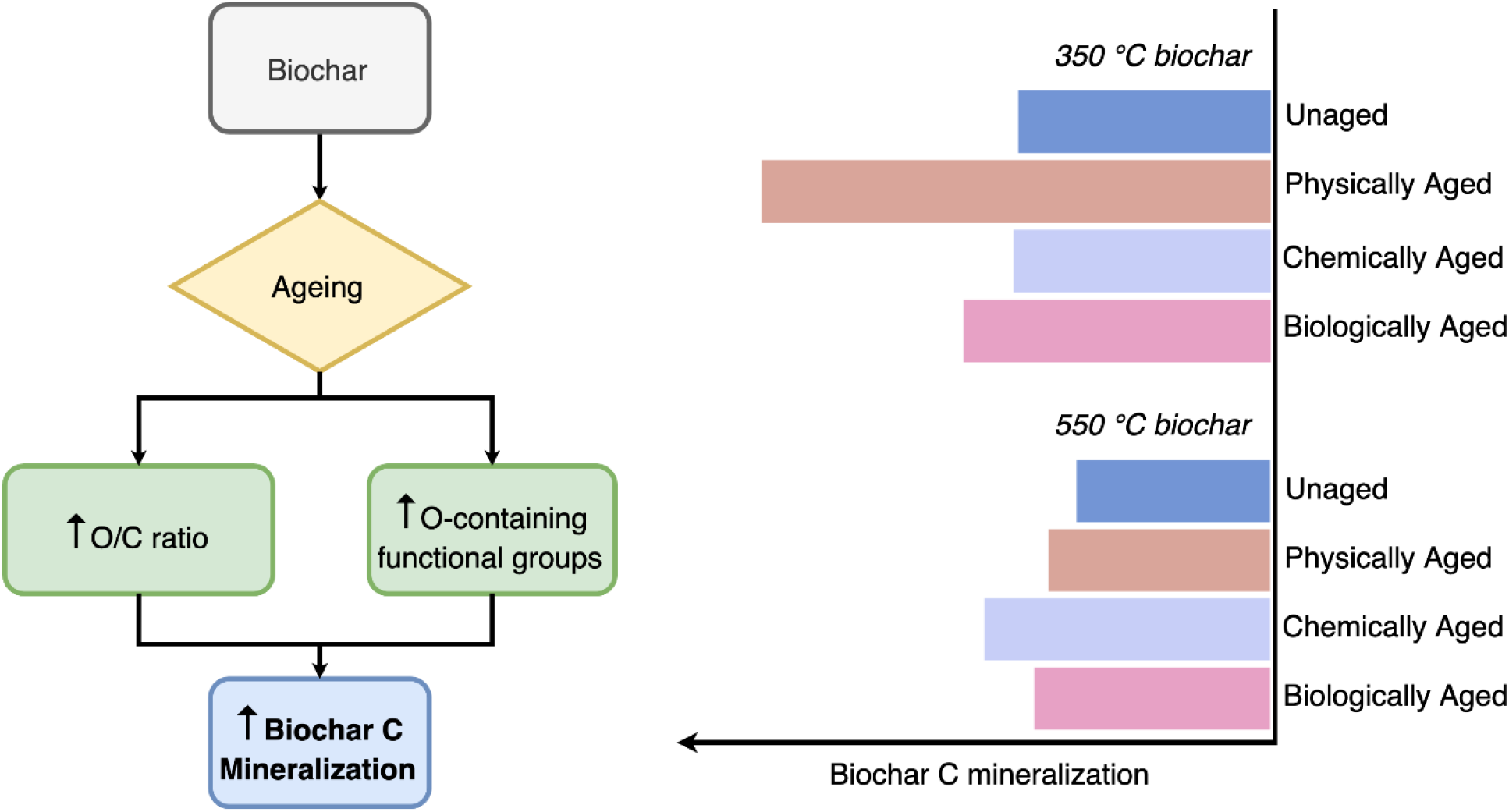

## 1. Introduction

Biochar is the carbon-rich solid product of pyrolysis, the process of heating biomass under oxygen limited conditions ^1^. Biochar has the potential to be used as a soil amendment for agricultural management (e.g., to increase water holding capacity, among other effects) and as a carbon (C) management strategy to help mitigate greenhouse gas emissions ^2^. Converting waste biomass into biochar can potentially be an effective way to sequester C, since the C contained in biochar is generally more resistant to mineralization compared to the C in the parent biomass ^2,3^, although the net C effects of biochar production and soil application depend heavily on system-specific parameters, particularly the baseline scenario for the fate of the parent biomass ^4–7^.

Biochar C is resistant to mineralization primarily because of its high proportion of condensed aromatic C ^2,8,9^, which has been shown to be resistant to mineralization by both abiotic and biotic processes ^10,11^. Further, biochar, while being rich in C, tends to have a low oxygen (O) content, and low O/C ratios in biochar have been shown to correlate with biochar persistence in soil ^12^. The chemical and physical properties of biochar that affect its persistence are initially determined by the production conditions, such as feedstock and production temperature ^13^, but once the biochar is deposited in soil, these properties change over time in a process known as ageing ^14–17^. Natural ageing of biochar in soil is a complex process ^14^, with multiple relevant mechanisms. We focus on three of these dominant mechanisms over the course of this paper:

1. *Physical ageing*-physical breakdown of biochar, primarily by freeze-thaw cycles and changes in temperature and moisture ^14,18–20^
2. *Chemical ageing*-degradation of biochar through abiotic oxidation upon exposure to various oxidizing agents ^21–23^
3. *Biological ageing*-biotic degradation and corresponding physical and chemical modifications of biochar by microbes and other soil organisms ^24–28^

Commonly reported effects of biochar ageing include a drop in pH, an increase in O content, and an increase in O-containing functional groups on the surface of aged biochar compared to unaged biochar. This suggests that ageing of biochar, both naturally and artificially, causes changes to its elemental composition and surface chemistry ^17^. Furthermore, these changes have been shown to affect properties of biochar such as sorption ^18,29^ and cation exchange capacity ^30,31^. However, we still have limited information on how these changes will affect soil CO_2_ emissions and the decomposability of biochar itself. Spokas ^32^ reported an increase in total C mineralization upon incubation of soil amended with 3 year aged woody biochar, primarily due to chemical oxidation of biochar surfaces. On the other hand, Liu *et al*. ^33^ observed lower total C mineralization in soil incubations amended with 6 year aged wheat straw biochar due to loss of easily mineralizable C during ageing. The specific effects of ageing on the decomposability of biochar by soil microbes have not been fully explored ^34^.

Further investigating the relationship between physicochemical changes and biochar decomposability is one of the primary tranches of this work. Specifically, the aim of this study is to examine the decomposability of aged biochar by a specific biochar decomposing microbe from a genus that is common to soils worldwide – a *Streptomyces* isolate. We predicted that the change in mineralization with ageing will depend on whether the ageing process results in loss of easily mineralizable C (as indicated by aliphatic chemical groups), which would lead to lower mineralization, or an increase in O content (as indicated by O/C ratios), which would lead to higher mineralization.

## 2. Materials and Methods

### 2.1. Production of biochar

Biochar was produced from eastern white pine wood chips (*Pinus strobus* (L.)) at highest treatment temperature (HTT) 350 and 550 °C in a modified Fischer Scientific Lindberg/Blue M Moldatherm box furnace (Thermo Fisher Scientific, Waltham, MA, USA) under continuous argon flow (1 L min^-1^) and a residence time of 30 min ^35^. Pyrolyzed material was ground using a ball mill and sieved to collect biochar with particle size <45 μm. The full details of biochar production can be found in Supplemental Note S1.

### 2.2. Ageing of biochar

Biochar produced at both 350 and 550 °C was subjected to one of three different ageing processes-physical, chemical and biological. We performed all ageing treatments on single batches of biochar to give us a final set of physically, chemically and biologically aged chars produced at 350 °C (350PHY, 350CHEM and 350BIO) and 550 °C (550PHY, 550CHEM and 550BIO). A batch of 350 °C unaged biochar (350UN) and 550 °C unaged biochar (550UN) acted as controls in our study.

#### Physical ageing

For physical ageing (PHY), we subjected biochar samples to 20 freeze-thaw-wet-dry cycles between −80 °C and 100 °C using pint-sized Mason jars (473.18 mL), building on the method reported by Hale *et al*. ^18^. We chose this wide temperature range based on findings by Cheng *et al*. ^14^ that demonstrated that ageing of biochar can occur over a temperature range from −22 °C to 70 °C and that greater ageing occurs at higher temperatures. Our method simulates an extreme scenario of severe weathering of biochar likely to occur across many seasonally snow-covered ecosystems in the northern hemisphere during precipitation-drying cycling and freeze-thaw cycling ^36,37^. Quartz sand (Sargent Welch, Buffalo Grove, IL, USA) was used to simulate a soil matrix (80 g with a 5% weight biochar amendment) and ultrapure water was added to the jars containing 4 g of biochar to achieve 40% water holding capacity (WHC). During each cycle, the jars were frozen at −80 °C for a median time of 7 hours (min 5 h – max 48 h), thawed for a period of 1-2 hours, following which they were dried in the oven at 100 °C for a median time of 18 hours (min 14 h – max 54 h) and cooled to room temperature for a period of 1-2 hours. After each drying period, masses of the jars were measured, and ultrapure water was added to reach 40% WHC. After 20 cycles, biochar particles were separated from the sand by wet sieving using a US mesh size no. 270 sieve that allowed the biochar particles less than 45 μm in size to pass through while retaining the sand particles.

#### Chemical ageing

For chemical ageing (CHEM), we treated biochar samples with H_2_O_2_ based on the method reported by Huff and Lee ^21^. We used a high concentration of H_2_O_2_ based on findings from previous studies that reported maximum changes in surface chemistry of biochar upon treatment with 30% w/w H_2_O_2_ solution ^21,38^. Briefly, 30% w/w H_2_O_2_ solution was added to 5 g of biochar at a ratio of 1 g biochar : 20 mL solution and shaken inside a chemical fume hood for 2 hours at 100 rpm. After 2 hours of shaking, we filtered the biochar samples through sterile Whatman glass microfiber filters (Grade 934-AH Circles – 1.5 μm particle retention) and rinsed with 100 mL aliquots of ultrapure water to remove any residual H_2_O_2_.

#### Biological ageing

For biological ageing (BIO), we exposed the biochar samples to a microbial community in a nutrient solution supplemented with glucose (40 μg glucose mg^-1^ biochar C), building on the method reported by Hale *et al*. ^18^. By adding glucose, we hoped to stimulate microbial activity and, with it, the decomposition of biochar. We chose a microbial community expected to be enriched in microbes that could degrade biochar to further accelerate the biological ageing treatment. We derived the microbial inoculum from soil samples collected at the Blodgett Forest Research Station at University of California, Berkeley, which has been used to conduct multiple prescribed burn studies ^39^. The soil samples for the inoculum were collected from 0-10 cm depth at the center of a slash pile burn after removing the ash layers. To extract the inoculum, we mixed the field-moist soil samples with Millipore water in sterile 50 mL centrifuge tubes and vortexed for 2 hours at high speed. After vortexing, the tubes were allowed to stand for 5 minutes and the soil suspensions were filtered through sterile 2.7 μm Whatman membrane filters into sterile centrifuge tubes. For the biological ageing process, nutrient solution was prepared from autoclave-sterilized modified basal salt solution (500 mL L^-1^ final biochar nutrient media), modified from Stevenson *et al*. ^40^, filter-sterilized vitamin B12 solution (200 μL L^-1^ final biochar nutrient media), filter-sterilized vitamin mixture (200 μL L^-1^ final biochar nutrient media) and a filter-sterilized trace element solution (1 mL L^-1^ final biochar nutrient media) ^41^. The detailed composition of the nutrient solution is provided as supplementary material accompanying this work (Supplemental Note S2.1.). We combined 5 g of biochar and glucose supplement (40 μg mg^-1^ biochar carbon) with 250 mL ultrapure water and autoclave-sterilized the mixture. After autoclaving, the biochar mixture was transferred to a quart-sized (946.35 mL) Mason jar and combined with 250 mL of the nutrient solution. Note that the pH of the modified basal salt solution, which is a part of the nutrient solution was adjusted to 7 to obtain a pH neutral final biochar nutrient media (Supplemental Note S2.1.). To the resulting biochar and glucose supplemented nutrient media, we added 8 mL of the filtered inoculum and incubated the jars at 30 °C in a shaker incubator set to 100 rpm for a period of 2 weeks.

### 2.3. Chemical Analyses

Total C and N were determined for aged and unaged biochar samples using a Thermo Scientific Flash EA 1112 Flash Combustion Analyzer (Thermo Fisher Scientific, Waltham, MA, USA) at the Department of Agronomy, UW-Madison, WI, USA. Total H was determined using a Thermo Delta V isotope ratio mass spectrometer interfaced to a Temperature Conversion Elemental Analyzer (Thermo Fisher Scientific, Waltham, MA, USA) at the Cornell Isotope Laboratory, NY, USA. Total O was calculated by subtraction as per Enders *et al*. ^13^, after determining ash content of aged and unaged biochar samples using the method prescribed by ASTM D1762-84 Standard Test Method for Chemical Analysis of Wood Charcoal (See further details in Supplemental Note S1).

The pH of aged and unaged biochar samples was measured in deionized water at a 1:20 solid: solution ratio using an Inlab Micro Combination pH electrode (Mettler Toledo, Columbus, OH, USA) connected to a Thermo Scientific Orion Star A111 benchtop pH meter (Thermo Fisher Scientific, Waltham, MA, USA). Further details of this procedure can be found in the Supplemental Note S1.

The FT-IR measurements were performed at the U.S. Dairy Forage Research Center, Madison, WI, USA with a Shimadzu IRPrestige-21 FT-IR spectrometer (Shimadzu, Kyoto, Japan) on the ATR (Attenuated Total Reflection) absorbance mode. Briefly, 5-10 mg of the biochar sample was placed on the Zn-Se sample trough and scanned. For each sample, we obtained 256 scans per sample in the range from 4000 to 650 cm^-1^ with a resolution of 1 cm^-1^ (550UN, 350PHY and 550PHY) and 2 cm^-1^ (350CHEM, 550CHEM, 350BIO, 550BIO and 350UN). Background corrections were performed between each sample measurement. We assigned wavenumbers for selected functional groups based on previous studies (S.I. Table S1) and quantified the peak heights of selected functional groups after spectrum normalization using the Shimadzu IR Solution FT-IR software (S.I. Table S2). Fractional signal heights for each of the FT-IR peaks were calculated by dividing the signal height of each of the peaks by the sum total of signal heights of all peaks of interest to determine the contribution of the signal generated by a particular species to the full spectra. Further details of this procedure can be found in the Supplemental Note S1.

#### Multivariate comparisons

To compare the full FT-IR spectra of biochar samples across temperatures and different ageing treatments, we used a multivariate dendrogram technique. We used the continuous normalized data for these analyses, excluding the region from 4000 cm^-1^-3100 cm^-1^ wavenumber to remove signals from water sorbed to the biochar surface. We used the dendextend ^42^ package in R to construct a dendrogram. Euclidean distances between biochar samples were calculated using the dist() function, and the hclust() function with the complete linkage method used for hierarchical clustering, where the two most similar samples are clustered together, one after another, forming an ordered hierarchical tree/ dendrogram.

### 2.4. Incubation

We performed the incubations with all the aged biochar (PHY, CHEM and BIO) as well as unaged biochar (UN) produced at both 350 and 550 °C as solid agar biochar media, inoculated with a bacterial isolate known to grow on biochar, while tracing CO_2_ emissions from each replicate.

The bacterial isolate we used was a *Streptomyces* that was isolated on media with eastern white pine wood biochar produced at 500 °C as the sole C source. The primary motivation for selecting this specific species is that it was able to grow on biochar media during trial lab incubations. Further, there is evidence that indicates that bacterial genera that respond positively to biochar addition in soils include members that have the potential to break down polycyclic aromatic hydrocarbons (PAHs) ^43^, a constituent of biochars, particularly high-temperature ones. We recovered the isolate from glycerol stocks by streaking onto a biochar (produced from pine wood at 350 °C) nutrient media agar plate (as described in Supplemental Note S2.) and incubating for 5 days at 37 °C. A single colony from the biochar media plate was inoculated into 30 g L^-1^ Tryptic soy broth (Neogen Culture Media, Lansing, MI, USA) and incubated at 30 °C in a shaking incubator until growth was visible, characterized by turbidity in the media.

We performed incubations in quarter-pint sized Mason jars (118.29 mL). Biochar (1 g L^-1^ final biochar nutrient media) and Noble agar suspension (30 g L^-1^ final biochar nutrient media) were sterilized by autoclaving and combined with nutrient solution to obtain a pH neutral final biochar nutrient media (Supplemental Note S2.). For each sample, we poured 40 mL of the final biochar nutrient media into sterile Mason jars. After the agar solidified, 20 μL of the bacterial suspension in malt extract broth was plated onto the agar surface using the spread plate technique ^44^.

We performed the incubations in replicates of at least three for each treatment and included uninoculated controls for each treatment. In addition, we included two empty jars as gas flux blanks for the experiment. After plating, the jars were capped and sealed with sterile, gas-tight lids provided with fittings that facilitate CO_2_ gas measurements and attached to randomly selected positions on the distribution manifolds (multiplexer) using polyurethane tubing ^45^. We measured the concentration of CO_2_ respired in the headspace of each jar at intervals of 3-4 days using a Picarro G2131i cavity ringdown spectrometer attached to the multiplexer over a period of 1 month. After each measurement, we flushed the jars with a 400 ppm CO_2_-air gas mixture to ensure aerobic conditions inside the jar. The precise concentration after flushing each jar was measured and subtracted from the next time point reading to determine the respired CO_2_ in the jar. From previous biochar incubation trials with the isolate, we confirmed that sampling over a 3-4-day interval did not lead to hypoxia inside the jars.

The raw CO_2_ readings measured using the multiplexer-Picarro system were processed in R to calculate biochar C mineralized over the period of incubation using the following packages: tidyverse ^46^, zoo ^47^, RColorBrewer ^48^, and broom ^49^. Briefly, we calculated the cumulative biochar C mineralized for each replicate at each time point. We corrected the cumulative biochar C mineralized values for all replicates by subtracting the corresponding mean C mineralized of uninoculated replicates within each treatment and a time series was plotted comparing the biochar C mineralization trends between the aged and unaged biochar samples.

#### Data Analysis / Statistical Methods

We performed most calculations in R using the packages dplyr ^50^ and tidyr” ^51^. Figures were made using the ggplot2 ^52^ and wesanderson ^53^ packages and all code used for analyses and figures in this paper is available at github.com/nayelazeba/biochar-ageing.

We performed ANOVA and Tukey’s HSD test in R to compare significant differences between cumulative biochar C mineralized across all aged and unaged biochar treatments for both 350 °C and 550 °C biochar. In order to correlate total C mineralized during the one-month incubation period across all aged and unaged biochar samples with fractional signal heights of each of the selected functional groups and molar O/C ratios, we performed linear regressions in R.

#### Microbial growth

After the incubation period, we disconnected the jars and analyzed images of the agar surfaces using the software *ImageJ* ^54^. The percentage area occupied by the growth of bacterial colonies was determined for each incubation jar and used as a rough proxy to compare microbial growth between jars (S.I. Fig. S1).

## 3 Results and Discussion

### 3.1. Effect of ageing on elemental composition

Aged biochars produced at both 350 °C and 550 °C had lower total C and higher total O contents than unaged biochar, except in the case of 350CHEM, where we did not observe similar trends (Table 1). The molar O/C ratio increased for 350BIO (0.26) and was highest for 350PHY (0.39) compared to 350UN (0.20) among 350 °C chars. In the case of 550 °C chars, the O/C ratio increased for 550BIO (0.18), 550PHY (0.15) and was highest for 550CHEM (0.26) compared to 550UN (0.11). This is consistent with previous studies that have shown an increase in O/C ratio following natural as well as artificial ageing of biochar through abiotic and biotic processes ^14,19,20,30,55^. The relative decrease in C with ageing is likely due in part to leaching of C-rich dissolved organic matter ^19^. Additionally, abiotic oxidation of C to carbon dioxide and utilization of C as a substrate by microbes in the case of biologically aged biochars is likely to result in relatively greater loss of C than O ^10,24,30,56^. The higher O content in aged biochars indicates an increase in O-containing functional groups that is likely due to both abiotic oxidation of C in the case of chemically and physically aged biochars ^14,30^ as well as microbially mediated oxidation in the case of biologically aged biochars ^55^. The effects of pyrolysis temperature on the elemental composition of biochar are discussed in Supplemental Note S4.

**Table 1.** Elemental Composition, Elemental Ratio, and pH of the Unaged and Physically, Chemically and Biologically aged biochar samples produced at low temperature (350°C) and high temperature (550 °C)^*^

| HTT<br>(°C) | Ageing<br>treatment | Total C | Total N | Total H | Ash | Derived<br>total O | O/C | pH in<br>solution |
| --- | --- | --- | --- | --- | --- | --- | --- | --- |
|  |  | (wt %) |  |  |  |  |  |  |
| 350 | Unaged | 75 ± 0.04 | 0.3 ± 0.00 | 3.9 ± 0.01 | 0.6 ± 0.02 | 20.4 | 0.20 | 6.1 ± 0.1 |
|  | Physical | 61 ± 1.03 | 0.3 ± 0.1 | 2.7 ± 0.04 | 4.3 ± 0.01 | 31.8 | 0.39 | 3.3<br>(N = 1) |
|  | Chemical | 80 ± 4.6<br>(N = 4) | 0.3 ± 0.04<br>(N = 4) | 2.5 ± 0.2 | 1.6 ± 0.04 | 15.6 | 0.15 | 4.3 ± 0.02 |
|  | Biological | 70 ± 1.2 | 0.3 ± 0.01 | 3.5 ± 0.2 | 1.8 ± 0.2 | 24.4 | 0.26 | 4.8 ± 0.06 |
| 550 | Unaged | 85 ± 1.2<br>(N = 4) | 0.2 ± 0.02<br>(N = 4) | 2.4 ± 0.2 | 0.8 ± 0.03 | 11.9 | 0.11 | 6.9 ± 0.08 |
|  | Physical | 79 ± 0.1 | 0.4 ± 0.02 | 2.2 ± 0.01 | 3.1 ± 0.2 | 15.8 | 0.15 | 6.5<br>(N = 1) |
|  | Chemical | 71 ± 0.2 | 0.3 ± 0.02 | 3.8 ± 0.4 | 0.7 ± 0.05 | 24.4 | 0.26 | 5.0 ± 0.2 |
|  | Biological | 77 ± 3.7 | 0.3 ± 0.01 | 2.5 ± 0.1 | 2.4 ± 0.1 | 18.2 | 0.18 | 4.8 ± 0.6<br>(N = 4) |

\*Note: Data shown represent the mean ± standard deviation based on duplicate measurements unless specified otherwise. While ageing caused changes in pH (See Supplemental Note S3 for discussion), we controlled for the effects of pH on microbial activity in this study by adjusting the pH of the final nutrient biochar media for all samples.

### 3.2. Effect of ageing on surface chemistry

Amongst the ageing treatments, physical ageing altered the surface chemistry of biochar the most (Fig. 1). For both 350 °C and 550 °C biochar samples, the spectra of physically aged biochar samples were the most dissimilar from the spectra of unaged biochar samples. While chemical and biological ageing also caused changes to surface chemistry, the effects appear to be less pronounced than in the case of physical ageing, as we see these samples cluster together more by production temperature than treatment method – *i*.*e*., production temperature was a more important determinant of biochar chemistry than ageing treatment. An important factor driving dissimilarity between 350 °C and 550 °C biochars is the increase in aromatic carbon content and decrease in H and O-containing functional groups on the surface of biochar with increasing pyrolysis temperatures ^10,11,57^ (See Supplemental Note S4 for further discussion on effects of pyrolysis temperature on surface chemistry). It is interesting to note that the two PHY aged samples are more similar to each other than to other samples produced at the same pyrolysis temperatures. This suggests that the physical ageing treatment had a stronger effect on the surface chemistry than production temperature.

**Figure 1.**
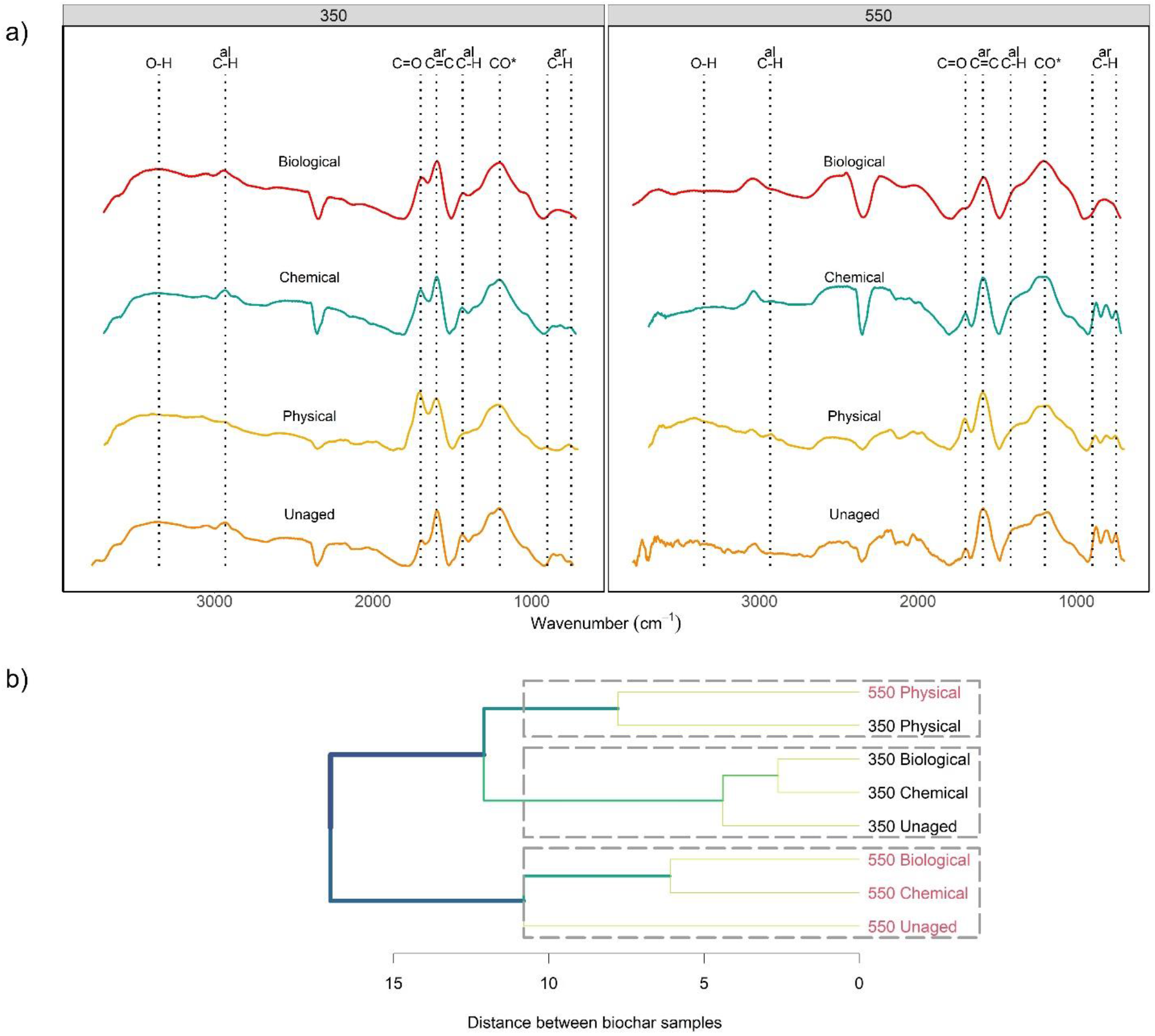
**(a)** FT-IR spectra of unaged and physically, chemically and biologically aged biochar samples produced at 350 °C (left panel) and 550 °C (right panel). Labels on top indicate the peak names assigned to different functional groups as described in detail in supplementary information (O-H: O–H stretching of carboxylic acids, phenols, alcohols at 3370 cm^-1^; al CH: aliphatic C-H stretch of CH_3_ and CH_2_ at ∼2932 cm^-1^ and C-H bending of CH_3_ and CH_2_ at 1413 cm^-1^; C=O: C=O stretch in carboxylic acids and ketones at ∼1701 cm^-1^; ar C=C: aromatic C=C vibrations and stretching of quinones at ∼1593 cm^-1^; CO*: C–O stretching, O–H bending of COOH and/or C–OH stretching of polysaccharides at ∼1200 cm^-1^; ar C-H: aromatic C-H out of plane deformation at 810 cm^-1^. **(b)** The clustering of biochar FT-IR spectra based on Ward’s hierarchical clustering method represented as a dendrogram. The distance of the link between any two clusters (or samples) is a measure of the relative dissimilarity between them.

#### Surface oxidation

An important feature that stood out when comparing the FTIR spectra of unaged and aged biochars was the increase in O-containing carboxylic groups, measured by changes in the relative peak height of the C=O stretch at 1701 cm^-1^ wavenumber (Fig. 1a and S.I. Table S2). Physically aged biochar across pyrolysis temperatures showed the maximum values for C=O stretch, indicating that the surfaces of physically aged biochar were the most oxidized and rich in carboxylic groups. Chemical ageing also resulted in surface oxidation. We measured a slight increase in carboxylic groups for both 350 °C and 550 °C chemically aged chars compared to unaged chars. The increase in surface oxygenation and O-containing functional groups after ageing is consistent with the findings of previous studies that investigated changes in surface chemistry using methods analogous to the physical and chemical treatments used in this study ^14,19–22^. For biological ageing, surface oxidation was observed only in the case of 350BIO. We did not observe any increase in the relative peak height of carboxylic groups in the case of 550BIO compared to 550UN. This suggests that abiotic oxidation through physical and chemical ageing methods used in the study resulted in more surface oxidation and carboxylic groups compared to biotic oxidation through biological ageing. This agrees with the finding of Cheng *et al*. ^30^, where they noted that abiotic processes were more important than biotic processes for the initial surface oxidation of fresh biochar. Our observations indicate that 350 °C biochars were more oxidized compared to 550 °C chars within a given ageing treatment (Fig. 1a and S.I. Table S2). This is most likely due to higher aromatic carbon content in biochar produced at 550 °C that tends to be more condensed and resistant to oxidation while the 350 °C chars have low aromatic carbon content that is less condensed and amorphous and more likely to undergo oxidation ^10,11^. This has been confirmed by other studies that observed an increase in resistance to oxidation by biochar produced at higher pyrolysis temperatures ^38,58,59^.

#### Surface aromatic and aliphatic groups

When considering how ageing affected the surface aromatic and aliphatic groups, we found that pyrolysis temperature was an important factor controlling these changes. For 350 °C biochars, we consistently observed a decrease in relative peak height in the 1413 cm^-1^ aliphatic C-H stretch, 810 cm^-1^ aromatic C-H stretch and 1593 cm^-1^ C=C aromatic stretch regions after ageing. The maximum decrease in peak values was consistently observed for 350PHY. Additionally, in the case of 350PHY, we measured a considerable decrease in relative peak height for the aliphatic C-H stretch at 2932 cm^-1^ after ageing. In the case of 550 °C char, we observed a considerable decrease in the relative peak height for the aromatic C-H stretch and a slight decrease in the 1413 cm^-1^ aliphatic C-H stretch after ageing but the same was not observed in the case of the C=C aromatic stretch. These changes indicate a relative loss or transformation of both surface aliphatic and aromatic carbon groups during ageing. As discussed earlier, the loss in C could be due to leaching or abiotic oxidation of C during ageing. Further, in the case of biological ageing, the relative loss in aliphatic C group at 1413 cm^-1^ and 2932 cm^-1^ (for 350°C chars) could be a result of decomposition of aliphatic C by soil microbes ^24,25,60^. While it may not be possible to conclusively determine whether oxidized functional groups were previously associated with aromatic vs. aliphatic compounds, the drop in relative heights in the aromatic regions (810 and 1593 cm^-1^ wavenumbers) accompanied by a relative increase in signal for carboxyl (1701 cm^-1^) group suggests that the oxidation of aromatic C results in the development of carboxylic groups. It has been previously suggested that oxidation on the edges of the aromatic backbone of biochar, taking place over a long period of time, could lead to the formation of negatively charged carboxyl groups ^56,61,62^. A loss in aromatic functional groups was documented during physical ageing of peanut straw biochar ^20^ and during chemical ageing of pine wood biochar ^21^. More recently, Yi *et al*. ^63^ measured loss and transformation of condensed aromatic C after 9 years of field ageing of high temperature bamboo and rice straw biochar. These previous findings support the inference that ageing methods used in this study could have caused the disruption of aromatic carbon to form carboxylic groups.

It is important to note that FTIR spectra as produced and analyzed in this study are only semi-quantitative – *i*.*e*., a doubling in peak height does not necessarily represent twice as much of the bond associated with that wavenumber. Furthermore, since replicates for ageing treatments and FTIR measurements were not included, we cannot determine whether these differences are statistically significant. However, the spectra represent an average of 265 scans on pooled and homogenized samples, and consistent responses to ageing at the two different temperatures as well as consistent temperature effects across different ageing treatments both help give us confidence in the trends observed here.

### 3.3. Effect of ageing on biochar C mineralization

The biochar C mineralization trends for all treatments show a similar pattern overall, with an initial period of steep increase in C mineralization, followed by the onset of a period of lower C mineralization (about 350 hours after the start of incubation, Fig. 2). This is comparable to the C mineralization curves commonly observed in previous incubation studies with soil amended with biochar ^28,33,64,65^ and biochar inoculated with a microbial community from a forest soil ^60,66,67^. Cumulative biochar C mineralized was significantly higher for 350 °C biochars compared to 550 °C biochars (Tukey test, p_adj_ = 0.003). This is consistent with previous studies that have noted higher microbial activity and respiration in incubations with low temperature chars ^56,67–69^, since biochar produced at high pyrolysis temperature contains a larger fraction of condensed aromatic C, which is more difficult for microorganisms to oxidize ^11,58,70–72^. Growth on agar surfaces over the month-long incubation, as measured by total surface area, was higher for 350 °C biochar treatments as compared to their 550 °C counterparts (S.I. Fig. S2). This corresponds to trends observed in the cumulative biochar C mineralized over the incubation period, indicating that the *Streptomyces* strain more effectively colonizes and grows on agar surfaces containing biochar particles produced at 350 °C.

**Figure 2.**
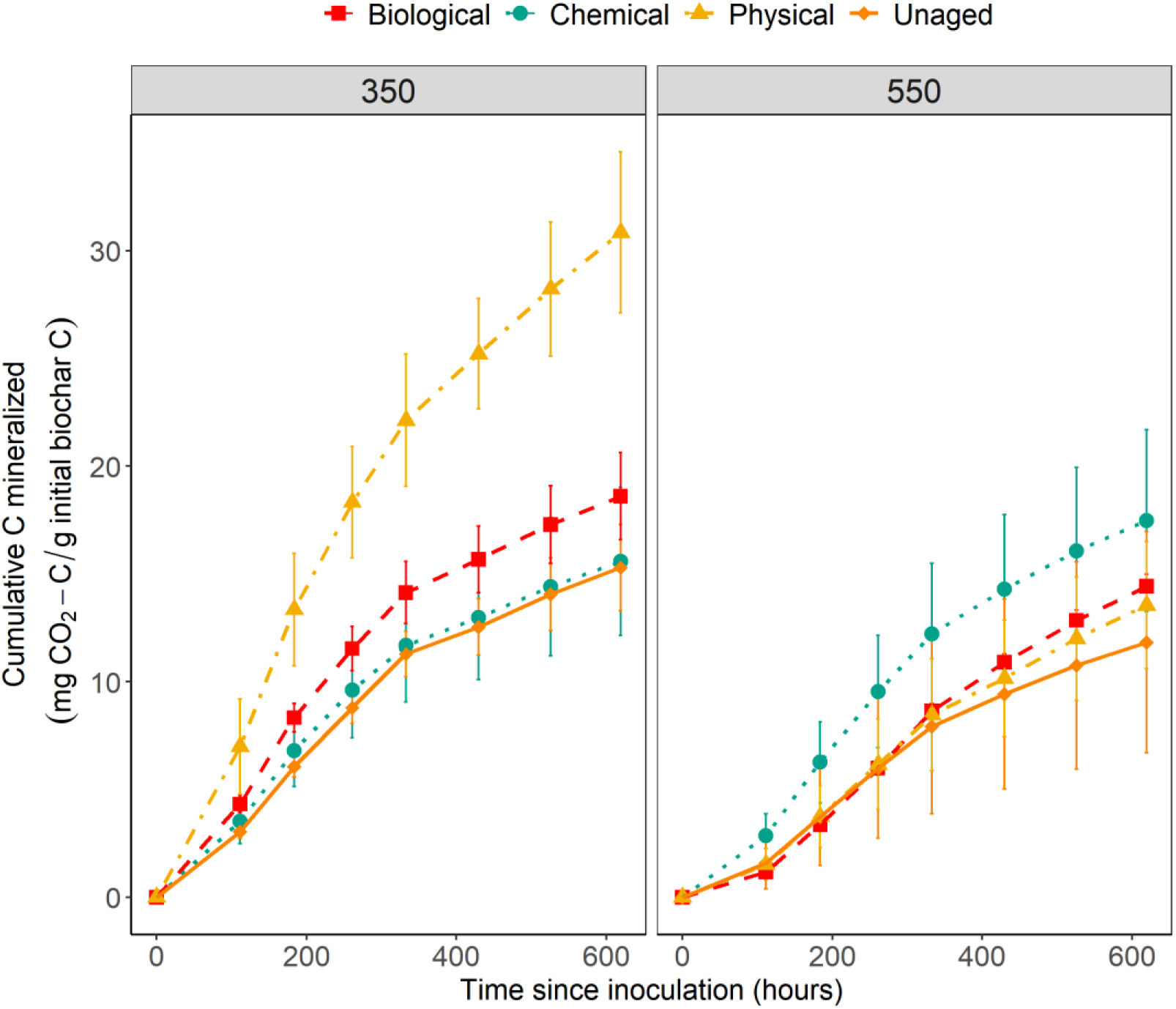
Mean cumulative C mineralized from unaged and physically, chemically and biologically aged biochar samples over time, with uninoculated blanks subtracted and normalized with mean biochar-C. N=3 for physical, chemical and unaged, N=5 for biological. Error bars represent 95% confidence intervals. The left panel shows biochars produced at 350 °C and the right panel shows biochars produced at 550 °C.

Amongst the 350 °C chars, the cumulative biochar C mineralized for aged biochars was higher than that for unaged biochar through the entire incubation period (Fig. 2), although the difference in C mineralization was statistically significant only in the case of 350PHY (Tukey test, p_adj_ = 0.0001). 350PHY treatments also showed the greatest surface growth (S.I. Fig. S2), consistent with the cumulative biochar C mineralized data.

Amongst the 550 °C biochars, there were not large differences between cumulative biochar C mineralization in aged versus unaged biochars. We observed an increase in biochar C mineralized for 550CHEM compared to unaged biochar through the incubation period and a slight increase in biochar C mineralization for 550PHY and 550BIO after about 400 hours after the start of incubation, but the differences in means were not significant (Fig. 2). These observations were consistent with trends in growth measurements on 550 °C biochar agar surfaces, where the average surface growth was 43% greater for 550CHEM compared to 550UN but no difference in growth was observed for 550BIO and 550PHY treatments (S.I. Fig. S2).

Biochar C mineralization across temperature and ageing treatments was significantly correlated with elemental composition (Fig. 3) and surface chemistry (Fig. 4). The greatest changes in surface and bulk chemistry were observed during physical ageing – specifically, we saw the greatest increase in O/C ratio and carboxyl groups and the greatest loss of aromatic and aliphatic C in 350PHY. These changes were accompanied by significantly higher biochar mineralization compared to unaged biochar. These correlations were notable across the full dataset for 350 °C chars. We identified a significant positive correlation between the O/C ratio and cumulative biochar C mineralized (R^2^=0.778, p<0.001; Fig. 3) and between relative peak height of carboxylic functional groups and cumulative biochar C mineralized (R^2^=0.731, p<0.001 for 1701 cm^-1^; R^2^=0.37, p=0.021 for 1200 cm^-1^; Fig. 4). Conversely, we identified a negative correlation between aromatic C and cumulative biochar C mineralized (R^2^ = 0.372, p=0.021 for 1593 cm^-1^; R^2^ = 0.464, p=0.0073; for 810 cm^-1^; Fig. 4) and between aliphatic C and cumulative biochar C mineralized (R^2^ = 0.869, p<0.001 for 1413 cm^-1^; R^2^ = 0.832, p<0.001; for 2932 cm^-1^; Fig. 4). These trends point to two factors that may primarily be responsible for the increase in mineralization with ageing for 350°C biochars:

**Figure 3.**
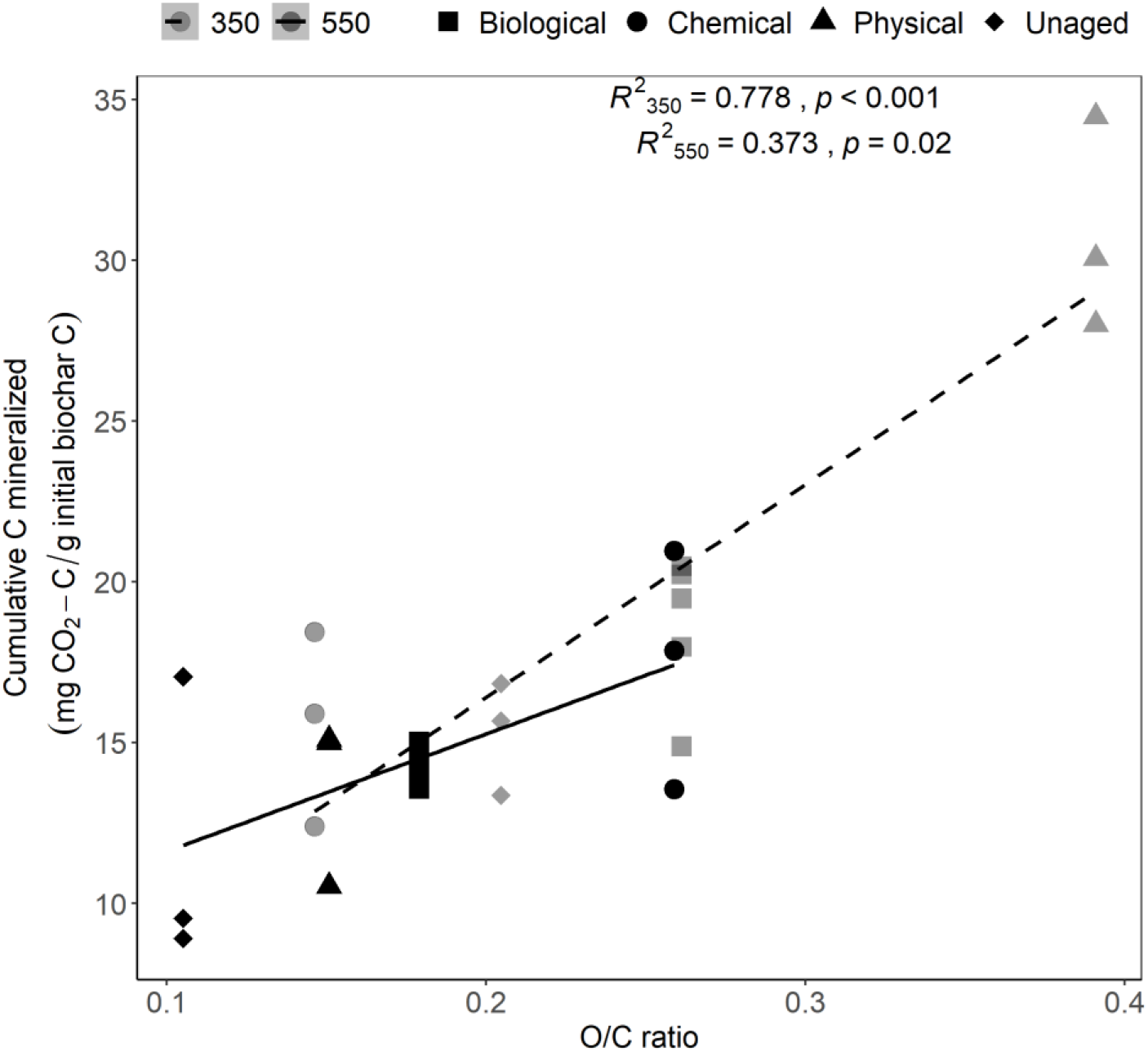
Relationship between cumulative biochar-C mineralized and molar O/C ratio. Shapes indicate unaged, physically, chemically and biologically aged biochar samples produced at 350 °C (light) and 550 °C (dark).

**Figure 4.**
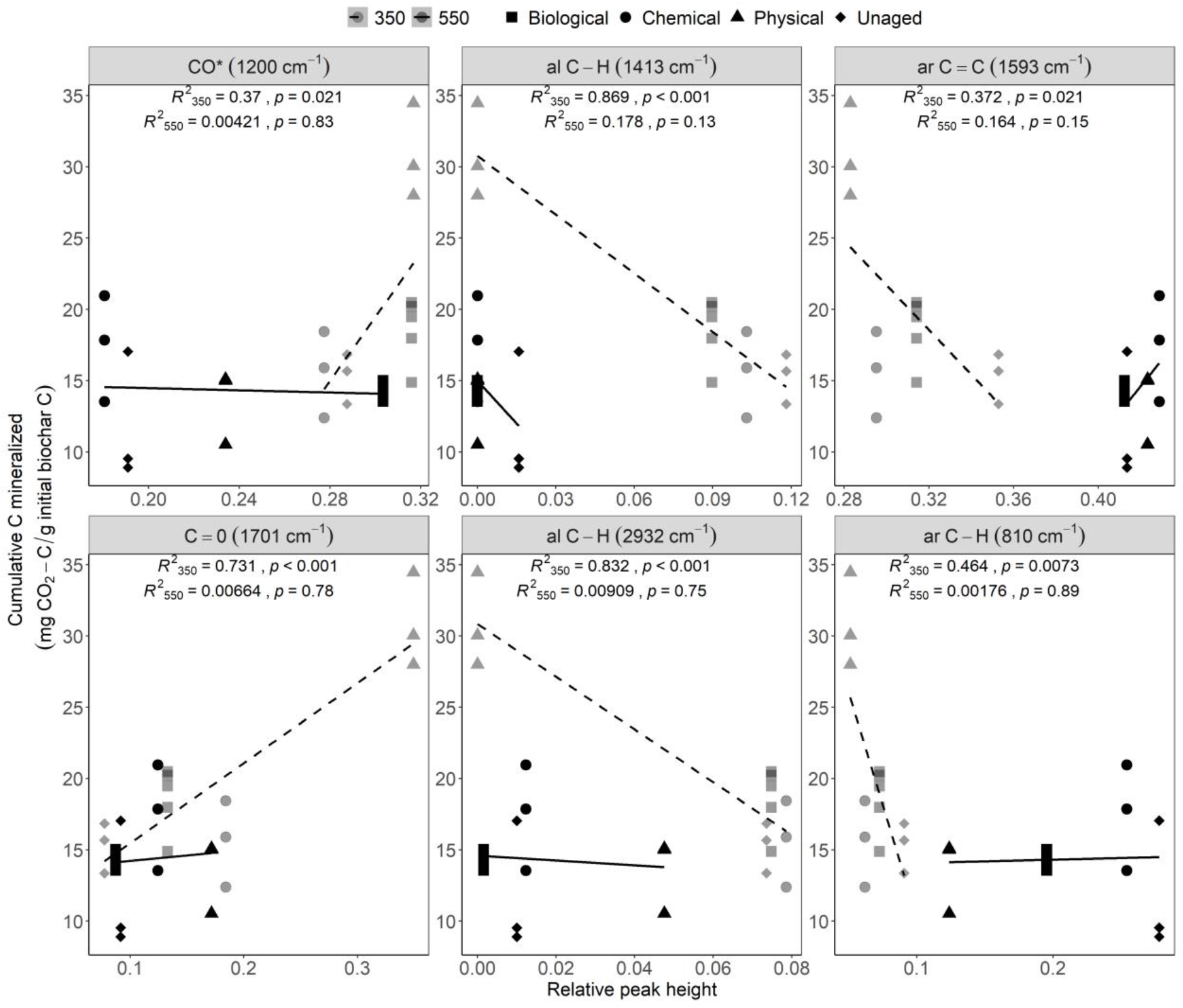
Relationship between cumulative biochar-C mineralized and FTIR spectra relative peak heights. Shapes indicate unaged, physically, chemically and biologically aged biochar samples produced at 350 °C (light) and 550 °C (dark). Panels indicate the peak names assigned to different functional groups at the given wavelength.

i. *higher O/C ratio of aged biochar:* An increase in carboxylic and phenolic groups during ageing could increase the O/C ratio of biochar, which makes it less stable, more hydrophilic and more likely to be mineralized by microbes ^12,66,72^. This surface-oxidized biochar is easier to break down and could potentially facilitate the microbial metabolism of ring structures that would ordinarily be highly recalcitrant ^14,17,56^.
ii. *oxidation/transformation of surface aromatic carbon to aliphatic C:* Oxidative transformation of aromatic C to linear alkyl-C and O-alkyl-C ^63^ could decrease ring condensation and make carbon more susceptible to microbial breakdown. Further, studies have documented breakdown and release of aromatic moieties in biochar to low molecular-weight organic acids during ageing ^73,74^.

For the 550 °C chars, we identified a positive correlation between the O/C ratio and cumulative biochar C mineralized (R^2^=0.373, p=0.02; Fig. 3) which suggests an increase in the mineralizability of biochar during ageing just as in the case of 350°C biochar. However, we were unable to identify any significant correlations between surface functional groups and cumulative biochar C mineralized (Fig. 4), which is perhaps to be expected, due to the non-significant differences in mineralization rates across 550 °C aged biochars. Even though increases in surface oxidation and loss of some surface aromatic and aliphatic C groups were observed during ageing, 550 °C aged chars still retained more aromatic C and were less oxidized than 350 °C aged chars as observed by measuring FTIR relative heights (S.I. Table S2). As a result, there is likely to be less easily mineralizable C present in 550 °C biochars, even after ageing. This is in line with other studies that have documented an inverse relationship between mineralization and aromatic fraction in biochars ^56,68^. This effect is potentially the primary reason we did not observe a significant increase in biochar C mineralized for 550 °C chars during ageing despite observing changes in the surface oxidation (Fig. 1a & Fig. 2).

This study provides evidence that higher O/C ratio and surface oxidation during ageing is likely to accelerate biochar C mineralization by microbes. In particular, the surface oxidation of the aromatic C groups has important implications for C management and cycling for both low temperature biochars and naturally produced wildfire pyrogenic organic matter (PyOM). It has been suggested that the chemical stability of natural wildfire PyOM produced at high temperatures is more comparable to low temperature biochars as natural PyOM was found to consist of small clusters of aromatic C units and not highly condensed polyaromatic structures ^75–77^. Based on our findings, the carbon in these PyOM materials could be more susceptible to surface oxidation and C loss during ageing which could lead to increased mineralizability and thus decreased C storage potential.

The ageing treatments used in this study were designed to simulate real world processes that occur naturally to biochars in soil. While it is not feasible to develop a scale to quantify the relative severity of our treatments compared to their expected severity in nature, our study highlights the role of abiotic factors like freeze-thaw-wet-dry processes in accelerating surface oxidation, which, in turn, increases the susceptibility of biochar to microbial degradation. There is a need to better understand the underlying mechanisms of surface oxidation in these processes and to design a more quantitative method to simulate ageing ^34^.

It is important to note that ageing and incubation of biochar was performed in the absence of soil (sand medium was used for physical ageing only), to control the processes of interest. However, we note that in soil systems, biochar-clay interactions, biochar-soil organic matter interactions as well as physical protection of biochar through aggregate formation are likely to affect both ageing of biochar and its interactions with microbes ^58,78,79^. Further investigation into changes in bulk and surface properties associated with long term ageing of biochar in biochar amended soils could help in verification of laboratory biochar ageing and incubation studies as well as broaden our understanding of the potential of biochar as a C sink ^34^.

## Supporting Information

FT-IR functional group peak assignments for biochar (Table S1); FTIR spectra relative peak heights (Table S2); Images of *Streptomyces* isolate growth on biochar-raw and processed using ImageJ (Figure S1); Comparison of *Streptomyces* isolate growth on aged and unaged biochar nutrient agar media (Figure S2); Details of biochar production and chemical analyses (Note S1); Details of biochar nutrient media preparation (Note S2); Effect of ageing on pH (Note S3); Elemental composition and surface chemistry of unaged biochar (Note S4).

## Author Contributions

The authors confirm contribution to the paper as follows: study conception and design: NZ, TLW; data collection: NZ, TDB; analysis and interpretation of results: NZ, TDB, TLW; draft manuscript preparation: NZ; manuscript review and editing: NZ, TDB, TLW. All authors have reviewed the results and have given approval to the final version of the manuscript.

## Notes

The authors declare no competing financial interest.

## Acknowledgements

This research was funded by the U.S. Department of Energy (DE-SC0020351; DE-SC0016365). We thank Akio Enders for supplying the biochar that we used to isolate *Streptomyces* and for assistance with biochar production. We are thankful to Kevin Panke-Buisse at USDA ARS, Maggie Phillips at the Jackson Lab and Kim Sparks at the Cornell Stable Isotope Laboratory for assistance with chemical analyses. Thanks to Jamie Woolet for isolating *Streptomyces* on biochar and for all the technical assistance with the incubations. We also thank Monika Fischer and Neem Patel for supplying the soil samples that we used during biological ageing.

