## Supplementary Material for "Effects of physical, chemical, and biological ageing on the mineralization of pine wood biochar by a *Streptomyces* isolate"

**Table S1.** FT-IR functional group peak assignment for biochar

| Functional group | Wavenumber ($cm^{-1}$) |
|---|---|
| O–H stretching of carboxylic acids, phenols, alcohols | 3370 [1,2] |
| Aliphatic C-H stretching | 2932 [1–3] |
| Carboxyl and carbonyl C=O stretching | 1701 [1–3] |
| Aromatic C=C vibrations | 1593 [1–3] |
| Aliphatic C-H deformation | 1413 [4,5] |
| *C–O stretching, O–H deformation of carboxylic groups and/or C–OH stretching of polysaccharides | 1200 [3,4] |
| Aromatic C-H out of plane deformation | 810 [6,7] |

*The peak at wavenumber 1200 $cm^{-1}$ detected could be a combination of peaks observed in the region 1260-1200 $cm^{-1}$: C–O stretching, O–H bending of COOH and 1170-950 $cm^{-1}$: C–OH stretching of polysaccharides.

**Table S2**. FTIR spectra relative peak heights

| Sample | ar C-H (810 $cm^{-1}$) | CO* (1200 $cm^{-1}$) | al C-H (1413 $cm^{-1}$) | ar C=C (1593 $cm^{-1}$) | C=O dbl (1701 $cm^{-1}$) | al C-H (2932 $cm^{-1}$) |
|---|---|---|---|---|---|---|
| 350 UN | 0.09 | 0.29 | 0.12 | 0.36 | 0.08 | 0.07 |
| 550 UN | 0.28 | 0.19 | 0.02 | 0.41 | 0.09 | 0.01 |
| 350 CHEM | 0.06 | 0.28 | 0.10 | 0.30 | 0.18 | 0.08 |
| 550 CHEM | 0.25 | 0.18 | 0.00 | 0.43 | 0.12 | 0.01 |
| 350 PHY | 0.05 | 0.32 | 0.00 | 0.28 | 0.35 | 0.00 |
| 550 PHY | 0.12 | 0.23 | 0.00 | 0.42 | 0.17 | 0.05 |
| 350 BIO | 0.07 | 0.32 | 0.09 | 0.31 | 0.13 | 0.07 |
| 550 BIO | 0.20 | 0.30 | 0.00 | 0.41 | 0.09 | 0.00 |

Note. Although the samples were dried before chemical analyses, we did not include the OH stretch peak at wavenumber 3370 $cm^{-1}$ in fractional signal heights calculations to avoid interference from any potential water molecules still sorbed to the biochar samples.

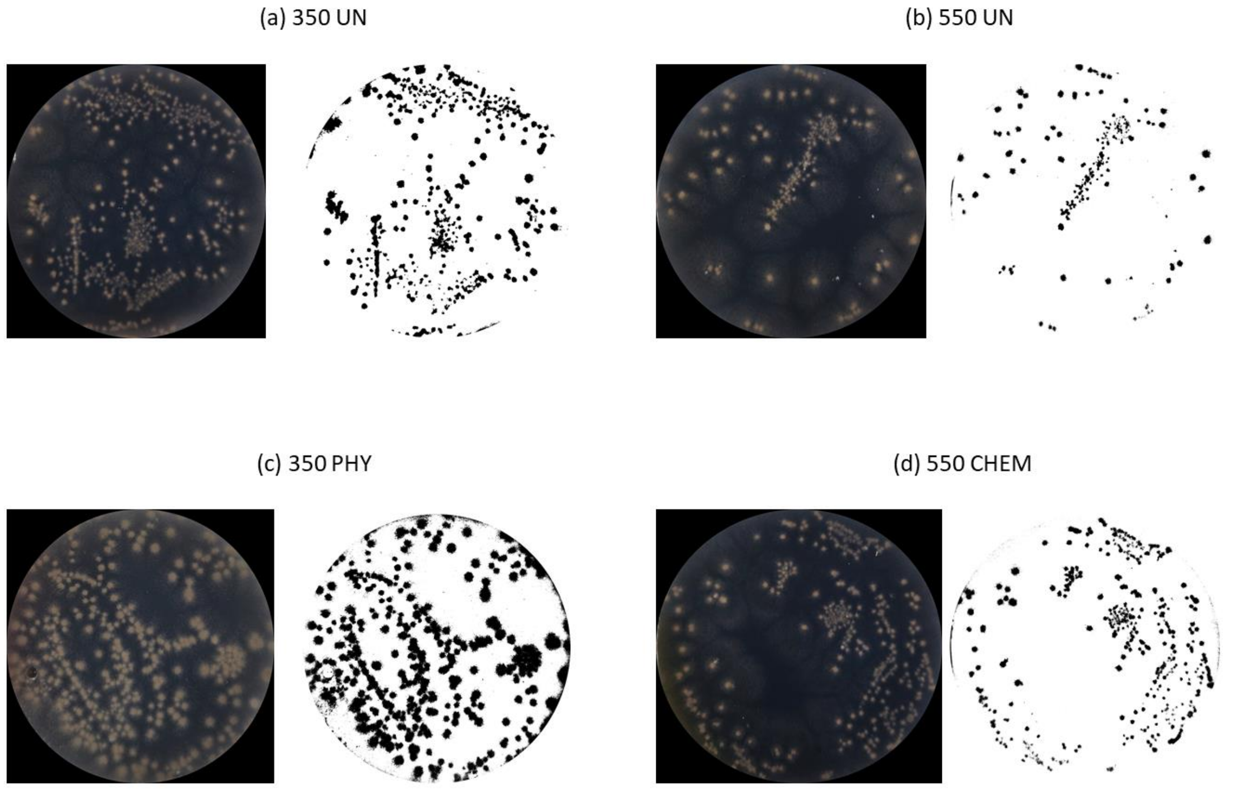


**Figure S1.** (Left) Images of *Streptomyces* isolate growth on the surface of biochar nutrient agar media at the end of the incubation period for a replicate of (a) 350 °C unaged biochar (b) 550 °C unaged biochar (c) 350 °C physically aged biochar (d) 550 °C chemically aged biochar. (Right) Processed images of *Streptomyces* isolate growth on the surface of the biochar sample shown on the left using the *Image J* software.

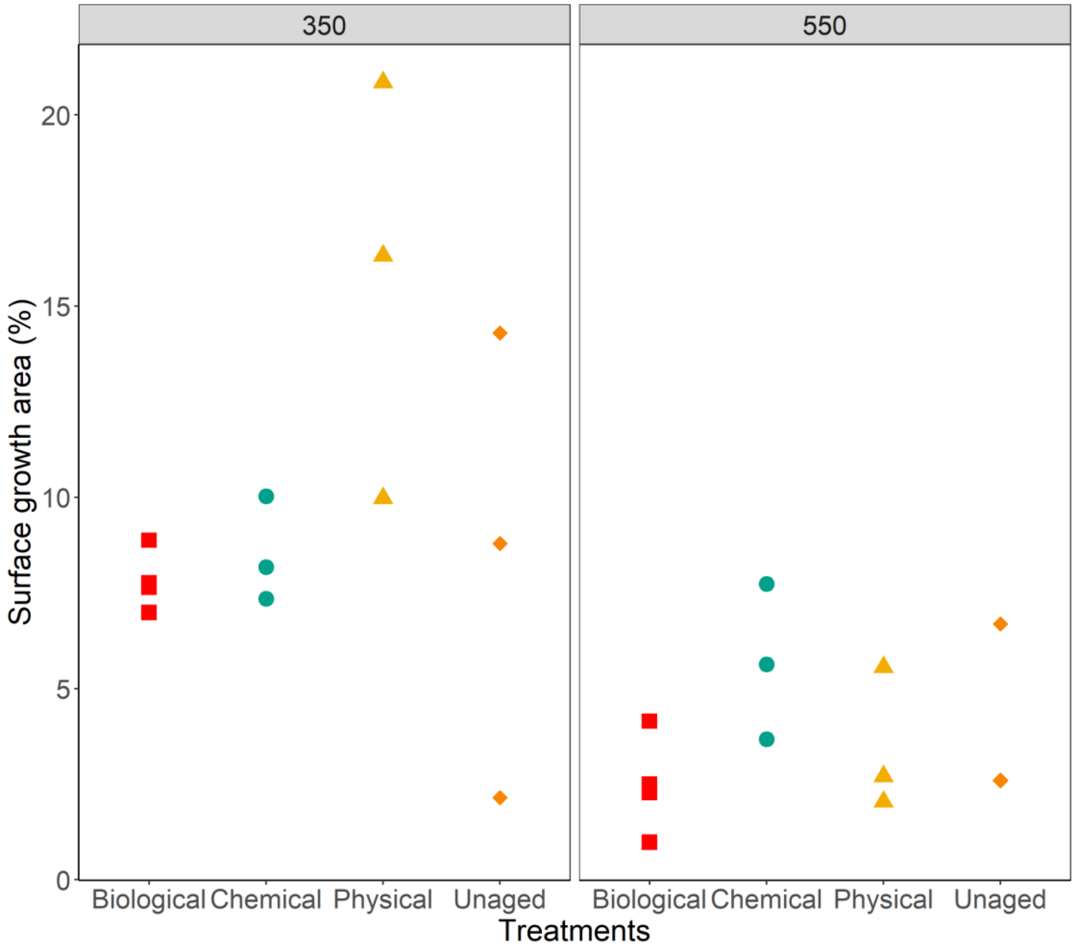


**Figure S2.** Growth of *Streptomyces* isolate on biologically, chemically, and physically, aged biochar and unaged biochar agar media over the incubation period. N=3 for physical, chemical and unaged, N=5 for biological. The left panel shows biochars produced at 350 °C and the right panel shows biochars produced at 550 °C.

### S1. Materials and Methods

*Production of biochar*

Biochar was produced from eastern white pine wood chips (Pinus strobus) at 350 and 550 °C in a modified Fischer Scientific Lindberg/Blue M Moldatherm box furnace (Thermo Fisher Scientific, Waltham, MA, USA) fitted with an Omega CN9600 SERIES Autotune Temperature Controller (Omega Engineering Inc., Norwalk, CT, USA) [8]. The feedstock was placed in a steel cylinder inside the furnace chamber and subjected to a continuous argon gas supply at a rate of 1 L $min^{-1}$ to maintain anaerobic conditions during pyrolysis. The heating rate for production of biochar was kept constant at 5 °C $min^{-1}$ and mixing of the feedstock inside the steel cylinder via mechanical paddles started once temperatures hit 250 °C. We held the temperature constant for 30 minutes once the highest treatment temperature was reached (350 and 550 °C), after which the product was rapidly cooled by circulating cold water in stainless steel tubes wrapped around the steel cylinder. Pyrolyzed material was ground using a ball mill and sieved to collect biochar with particle size <45 µm.

*Chemical Analyses*

Elemental analysis: Total C and N were determined for aged and unaged biochar samples using a Thermo Scientific Flash EA 1112 Flash Combustion Analyzer (Thermo Fisher Scientific, Waltham, MA, USA) at the Department of Agronomy, UW- Madison, WI, USA. Total H was determined using a Thermo Delta V isotope ratio mass spectrometer interfaced to a Temperature Conversion Elemental Analyzer (Thermo Fisher Scientific, Waltham, MA, USA) at the Cornell Isotope Laboratory, NY, USA. In order to estimate total O by subtraction, we determined ash content of aged and unaged biochar samples using the method prescribed by ASTM D1762-84.

Replicates were included for all samples for the CHN elemental analysis and ash measurements. Total O was calculated by subtraction as per Enders *et al.* [9] as follows:

O (%w/w) = 100 - C (%w/w) - N (%w/w) - H (%w/w) - ash (%w/w)

pH: The pH of aged and unaged biochar samples was measured in deionized water at a 1:20 solid: solution ratio using an Inlab Micro Combination pH electrode (Mettler Toledo, Columbus, OH, USA) connected to a Thermo Scientific Orion Star A111 benchtop pH meter (Thermo Fisher Scientific, Waltham, MA, USA). Briefly, we added 0.15 g of biochar to 3 mL of deionized water in 5 mL centrifuge tubes and vortexed for 1 hour at low speed. After vortexing, the tubes were centrifuged at 15000 X g for 2 min to sediment the biochar particles. We aliquoted 200 μL of the clear suspension into microtubes and measured the pH.

Fourier-transform infrared (FT-IR) spectroscopy: We quantified the heights of selected functional groups were quantified using the Shimadzu IR Solution FT-IR software. The spectra were ATR corrected to approximate a transmission spectrum and smoothed to reduce the background noise. We drew baselines for the spectra in the following regions using a quartic fitting function: 3800-2300 $cm^{-1}$, 2300-1820 $cm^{-1}$, 1820-1500 $cm^{-1}$, 1500-925 $cm^{-1}$ and 925-700 $cm^{-1}$. Wavenumbers were assigned for selected functional groups based on studies as described in Table S1. Peaks were manually detected around specific wavenumbers by identifying peak maxima, upper and lower baselines for each of the functional groups. Baselines for selected peaks were drawn as follows: 3006-2783 $cm^{-1}$ for aliphatic C-H stretch, 1808-1645 $cm^{-1}$ for C=O stretch, 1670-1483 $cm^{-1}$ for C=C vibrations and stretch, 1501-1307 $cm^{-1}$ for C-H bending of CH2 and CH3, 1334-919 $cm^{-1}$ for C–O stretching, O–H bending of COOH and/or C–OH stretching of polysaccharides and 934-713 $cm^{-1}$ for aromatic C-H deformation.

**S2. Biochar nutrient media preparation**

The final biochar nutrient media used for biochar incubations is a combination of the following components:

Nutrient solution (described in detail in S2.1.), which is composed of:

Modified basal salt solution

Trace elements solution

Vitamin mixture

Vitamin B12

Agar and biochar suspension

<u>To prepare the final biochar nutrient media agar plates (per 1 L):</u>

Agar and biochar suspension = 500 mL

Modified basal salt solution = 500 mL

Vitamin B12 = 200 uL of 250 mg L-1 solution* , **

Vitamin mix = 200 uL* , **

Trace elements solution = 1 mL* , **

*To be added after autoclaving

**Filter sterilize with 0.2 micron filter

<u>To prepare biochar agar suspension (per 1 L):</u>

Nobel agar = 60 g

Ground biochar = 2 g

**S2.1. Nutrient solution**

Basal salt solution (per 1 L):

$KH_2PO_4$ = 0.4 g

$NH_4Cl$ = 0.5 g

KCl = 1.0 g

$CaCl_2$. $2H_2O$ = 0.3 g

NaCl = 2 g

$MgCl_2$. $6H_2O$ = 1.24 g

$NaSO_4$ = 5.86 g

This solution will end up being acidic with a pH of ~6, The pH was adjusted to 7 by addition of NaoH and buffered by adding 1 g of MES (2-(N-Morpholino) ethane sulfonic acid) (VWR International, Radnor, PA, USA) before autoclaving.

Trace Elements Solution (aka SL-10, per 1 L):

HCl 25% (v/v) = 10 mL

$FeCl_2$. $4H_20$ = 1.5 g

$CoCl_2$. $6H_20$ = 190 mg

$MnCl_2$. $4H_20$ = 100 mg

$ZnCl_2$ = 70 mg

$H_3BO_3$ = 6 mg

$Na_2MoO_4$. $2H_2O$ = 36 mg

$NiCl_2$. $6H_2O$ = 24 mg

$CuCl_2$. $2H_2O$ = 2 mg

Vitamin Mixture (per 1L):

4-aminobenzoic acid (Vitamin B9 precursor) = 40 mg

D(+)-biotin (Vitamin B7) = 10 mg

Nicotinamide (Vitaman B3) = 100 mg

D(+)-pantothetic acid hemicalcium (Vitamin B5) = 50 mg

Pyridoxamine dihydrocloride (Vitamin B6) = 100 mg

Thiamine dihydrochloride (Vitamin B1) = 100 mg

### S3. Effect of ageing on pH

For unaged biochar, the pH of 550UN was higher (mean = 6.9) than the pH of 350UN (mean = 6.1; Table 1). This is consistent with previous studies that reported an increase in the pH of pine biochar with increasing pyrolysis temperatures [9–11]. The increase in pH is attributed to the reduction of organic functional groups, such as –COOH and –OH at higher temperatures [9,11]. Another factor that contributes to the rise in pH at higher pyrolysis temperatures is the relative increase of ash content in the biochar, although this effect is smaller in low ash containing woody biochars [9].

We observed a decrease in the pH of aged biochar compared to unaged biochar both at 350 °C and 550 °C (Table 1). We expect that the methods used in the preparation of the samples are primarily responsible for the change in pH, largely through the leaching of ash as all the treatments were prepared in a liquid medium. Additionally, leaching could have occurred during wet sieving to separate aged biochar particles in the physical ageing treatment and during rinsing and filtering of aged biochar particles in chemical and biological ageing treatments. In addition to the loss of ash during ageing, for the physically and chemically aged biochars, the decrease in pH may also be due to an increase in carboxyl functional groups for chemically and physically aged biochars when compared to unaged biochars for both 350°C and 550°C chars (Fig. 1a; See further discussion in section 3.3 in the main article). This is consistent with the findings of previous studies with similar chemical [12] and physical [13] ageing treatments. In contrast to the physically and chemically aged biochars, it is unclear if the decrease in pH of BIO-aged biochar could also have been partly a result of an increase in the amount of carboxyl functional groups on the surface. We were unable to identify any changes in the surface chemistry of biologically aged biochars compared to unaged biochars through FT-IR measurements (Fig 1a). Thus, we expect that leaching of ash is primarily responsible for the decrease in pH of BIO-aged biochars.

### S4. Elemental composition and surface chemistry of unaged biochar

*Elemental analysis*

For unaged biochar, the total C in 550UN was higher (mean = 84.7%) than that in 350UN (mean = 74.8%), while the total O and H contents were lower in 550UN (mean O = 11.9%; mean H = 2.4%) compared to 350UN (mean O = 20.4%; mean H = 3.9%; Table 1). This is consistent with previous studies that have reported an increase in the total carbon content of biochars with increasing pyrolysis temperature, due to increased carbonization and greater relative loss of H and O [9,11].

*FTIR*

While comparing the individual FTIR peaks between 350UN and 550UN, we observed changes in regions associated with aliphatic and aromatic C groups (Fig. 1a). There was, however, no notable change in relative height for the C=O stretch peak in carboxylic acids from 350UN (0.08) to 550UN (0.09). For the aromatic C functional groups, we observed an increase in the relative peak height with increasing pyrolysis temperatures (Fig. 1a and Table S2). The greatest increase appeared in the 810 $cm^{-1}$ aromatic C-H out of plane deformation (0.09 to 0.28) and a slight increase was seen in the 1593 $cm^{-1}$ C=C aromatic stretch (0.36 to 0.41). In contrast, the relative peak heights of groups indicative of aliphatic and lignin/cellulose-derived transformation products that are present in low temperature biochar decreased with increasing pyrolysis temperature. We observed a slight decrease in the relative peak height from 350UN to 550UN for 2932 $cm^{-1}$ aliphatic C-H stretch of $CH_3$ and $CH_2$ (0.07 to 0.01), 1413 $cm^{-1}$ C-H bending of $CH_3$ and $CH_2$ (0.12 to 0.02) and 1200 $cm^{-1}$ C–O stretching of phenols/ COOH/ polysaccharides (0.29 to 0.19). With the exception of the C=O carboxyl stretch, these broad trends in surface chemistry with increasing pyrolysis temperature are consistent with previous studies that investigated the effect of pyrolysis temperature on the chemical properties of biochar [14–16].
